# The carbon footprint of science when it fails to self-correct

**DOI:** 10.1101/2025.04.18.649468

**Authors:** Martin Farley, Marcus R Munafò, Anna Lewis, Benoit P. Nicolet

## Abstract

Science is – in principle – self-correcting, but there is growing evidence that such self-correction can be slow, and that spurious findings continue to drive research activity that is no longer justified. Here we highlight the environmental impact of this failure to self-correct sufficiently rapidly. We identified a non-fraudulent occurrence of irreproducible findings: the literature on the association between genetic variation in the serotonin transporter gene (5-HTTLPR) with anxiety and depression. An initial report in 1996 found evidence for an association, but a study as early as 2005 that was three orders of magnitude larger found no evidence for an association. However, studies investigating this association continue to be published. We isolated 1,183 studies published between 1996 and 2024 that investigated the association and calculated an estimated carbon footprint of these studies. We estimate that the failure to self-correct had a footprint of approximately 30,068 tons of CO2 equivalent. Our aim is to present a case study of the potential carbon footprint of research activity that is no longer justified, when a theory is disproven. We highlight the importance of integrating self-correction mechanisms within research, and embracing the need to discontinue unfruitful lines of enquiry.

## INTRODUCTION

Science self-corrects, in principle, but the speed and efficiency with which it does this is an empirical question^1^. It is inevitable that the scientific process will generate false positive findings, but self-correction broadly requires three things to happen^2^. First, researchers must attempt to replicate those findings. Second, those replication attempts must be analysed and interpreted through a neutral, disinterested lens and reported transparently. And third, if evidence accumulates that the original finding was erroneous then the community must admit the error, and potentially terminate that line of enquiry, in order to move on to more fruitful questions. Unfortunately, academic researchers are under significant pressure to publish findings, validation studies are often not prioritized, and any well-supported challenges to a given hypothesis are often harder to publish than if conforming to existing line of thoughts. Together, these factors can act against all three aspects of scientific self-correction.

An example of the failure to self-correct is the association between a length polymorphism of the serotonin transporter gene (5-HTTLPR) and anxiety related traits. In a 1996 paper, the “short” version of this polymorphism was reported to be associated with traits of anxiety^3^. Subsequent studies provided a putative mechanism – amygdala response to threat in brain imaging studies^4^ – and suggested that gene-environment interplay may play a part, with the “short” allele leading to a stronger response to stressful life events, as reflected in an increased risk of depression^5^. The combination of these lines of evidence built a compelling and elegant narrative – just, unfortunately, one that was repeatedly disproven^6,7,8^.

It is difficult to pinpoint when the tide of evidence turned. However, by 2009 large primary studies and meta-analyses – including of the putative moderating role of 5-HTTLPR on the effect of stressful life events on depression – had provided evidence that the apparent association was in fact artefactual ^6,7^. Around the same time, the study of genetic associations with complex behavioural phenotypes moved on from “candidate gene” approaches (where specific genetic variants of *a priori* interest were studied) to whole genome approaches (i.e., genome-wide association studies that agnostically scan the entire genome in very large samples). Notably, whole genome approaches proved to be remarkably efficient at identifying associations with a range of phenotypes that replicated reliably, but did *not* find evidence for a role of the serotonin transporter gene polymorphism in depression or anxiety. Subsequent large-scale investigations confirmed the lack of association of 5-HTTLPR genetic polymorphism with these outcomes^8^.

One might have expected the door to close on the 5-HTTLPR literature; unfortunately, it did not. Although the rate of publication has declined since 2013-2015, a PubMed search for “5-HTTLPR” will reveal recent publication (of both claimed findings and null results), adding to a field that can now be considered an exercise in noise generation. The impact of this is not negligible and adds extra strain on the academic system. Studies require research time and resources, while additional manuscripts add stress onto editorial boards and reviewers.

Scientific research is resource intensive^9^. Any research that takes place within a laboratory environment will require significant investments in infrastructure, energy, consumables, travel, time and money. A laboratory facility can consume ten-times more energy than a comparable office space, largely due to increased ventilation and equipment needs^10^.

Energy consumption is categorised as Scope 1 and Scope 2 emissions. The former represents direct emissions linked to combustion, typically from gas boilers in laboratory settings to maintain specific environmental conditions necessary for research. Electricity, falling under Scope 2 emissions when sourced from an external supplier, is consumed by plug-load equipment, air handling units, and other plant equipment. Manufacturing, usage and disposal of single-use consumables, chemicals and reagents accumulates to even greater environmental impacts of laboratory research. These, alongside travel, make up Scope 3 emissions, typically constitute the largest proportion of carbon emissions for any organisation or research context^11^. Impacts are further exacerbated by frequent travel either due to commuting or to communicate and engage the research community with findings. As long as incentives to produce more publications at high volumes remain, there will be an associated consumption and potential waste of resources associated with the generation of findings regardless of the capacity for self-correction.

Today, investments in science and research stand at nearly 2% of the global GDP^12^, and while its full impacts on the environment have yet to be fully quantified, case studies show that it is having an increasing demand on our global environment as well^13^. As we stand in the midst of a climate crisis, science has an intrinsic responsibility not only to assist in addressing global challenges, but also ensuring it does not excessively contribute to them. Here, we attempt to estimate the environmental and financial cost of the failure of the 5-HTTLPR literature to self-correct. Our findings highlight the importance for self-correcting mechanisms to ensure the reproducibility of findings, and spare financial and environmental costs.

## RESULTS

We first isolated 1,183 publications from 1996 to July 2024 linking 5-HTTLPR with anxiety or depression (excluding reviews, see methods). Studies disproving a role for 5-HTTLPR appeared in 2005^6^ and 2009^7^ (Figure 1A). Using these dates, we counted the research articles published after these two studies. We found that 1,052 articles 89% of total were published between 2005 and 2022, and 867 (73% of total) articles between 2009 and 2022. Even when factoring in a delay in information spreading after the 2009 paper, of 1 or 2 years, 779 (66% of total) and 704 studies (60% of total) were published after the respective delays. This indicates that the majority of articles on 5-HTTLPR appeared after the publication of strong evidence disconfirming a meaningful role of 5-HTTLPR in anxiety and depression.

**Figure 1.**
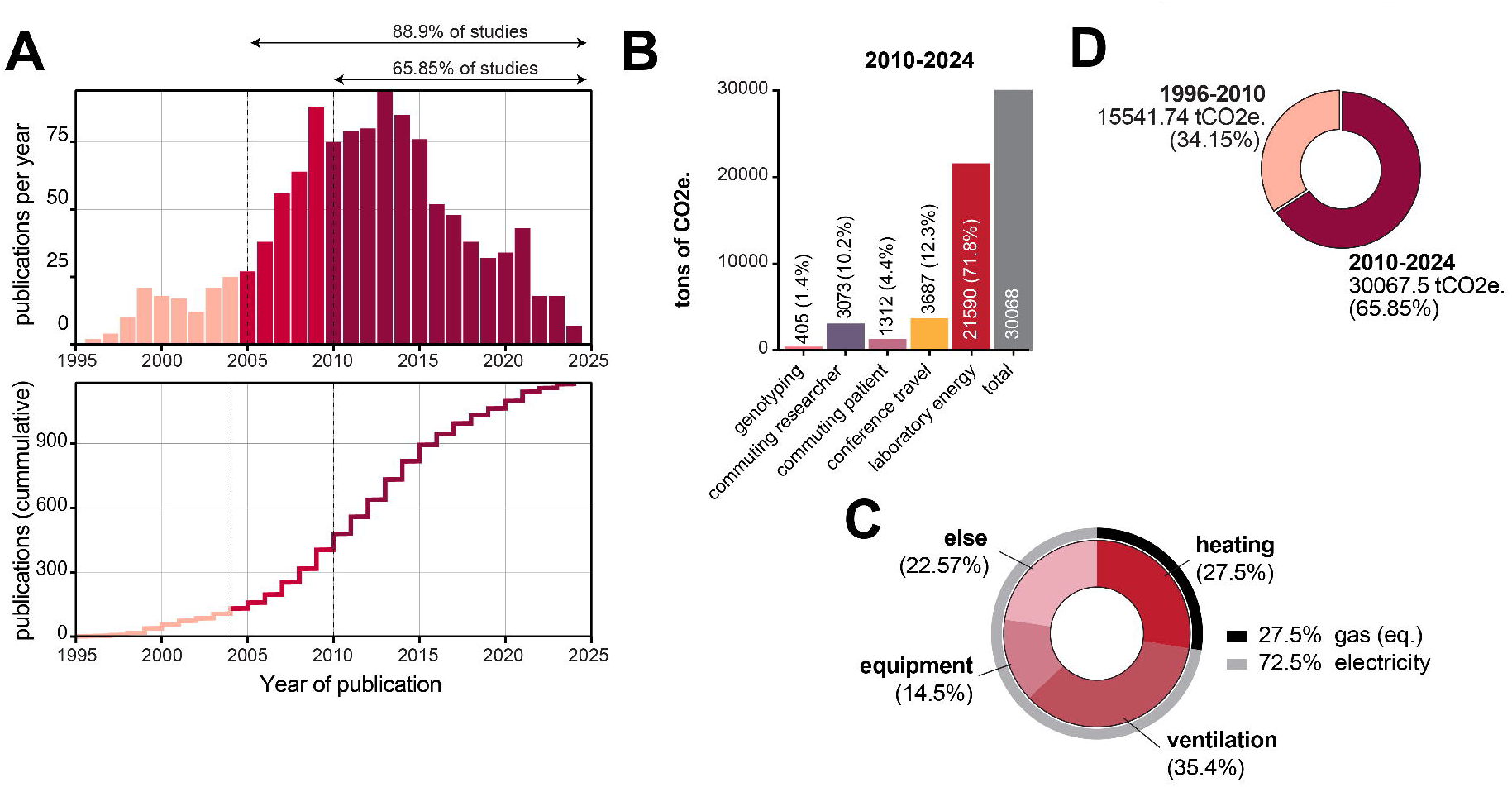

Beyond the moral implication of a failure of a field of research to correct itself, there is also a significant financial and environmental cost. To estimate the CO2e footprint of the 779 studies published between 2010 and 2024 (leaving >5 years for the field to adapt from the 1^st^ publication refuting the findings), we assessed the impact of four key areas: genotyping, commuting, conference travel, and laboratory building energy consumption (see methods).

### Genotyping Carbon Footprint

We first focused on the CO2e footprint estimation of genotyping. With 755 genotype assays per study on average^18^, a total of 893,165 genotyping assays were estimated to have been performed, amounting to 616.3 tons of CO2e for all the 1,183 studies. For all studies published from 2010 to 2024, we estimated that 588,145 genotyping assays have been performed totalling to 405.8 tons of CO2e (Figure 1B). Our estimates are likely to be an underestimation as we assume 100% success rate per genotyping reaction.

### Commuting Carbon Footprint

Next, we estimated the CO2e footprint of commuting. We employed published estimates of commute (US), where on average, one-way commutes amounted to approximately 13 miles, or nearly 21 km^19^. For all studies, we estimated that footprint from commuting ranged from 2,308 to 13,846 tons of CO2e assuming 0.5 to 3 years, respectively, of full-time work from start to completion of work (Figure S1). When sub-setting these estimates to studies published between 2010 and 2024, commuting amounted to between 1,537 (0.5y) and 9,219 (3y) tons of CO2e (Figure S1; see methods). Of note, we do not take the commuting of patient to the care centre for sample collection which would amount to 1,992.2 tons of CO2e (1996-2024) and to 1,311.8 tons of CO2e (2010-2024; Figure 1B). The mode of transportation (bike, train, car) will influence the overall footprint of commuting. Together, this suggests that the estimated footprint of commuting is greater than that of genotyping.

### Conference Travel Carbon Footprint

In recent years, studies have highlighted the environmental costs of conferences^14,15,16^. Using existing data on the travel of scientists based on job title^17^, we estimated that each author resulted in 1,057 kg CO_2_ e per annum. According to empirical data^17^, academic travel 2.2 times a year per person. Assuming 2 authors travelling per conference, we estimate 5,599 tons of CO2e (1996-2024) and to 3,687 tons of CO2e (2010-2024; Figure 1B). We likely underestimated the total footprint of conferencing as we only accounted for travelling^18,19^. The estimated footprint of conferencing is greater than both genotyping and/or commuting.

### Laboratory Energy Carbon Footprint

Finally, we assessed the footprint of laboratory energy consumption including electrical and heating/cooling demand. To assess the surface of laboratory space needed be considered, we used the number of authors as a proxy for lab size, and derived the surface per lab from a university report. The author numbers per study is 7.4 (mean) and 7 (median) ranging from 1 to 93 authors. While all 7 authors are unlikely to work full time for the study, the operational energy demands are left virtually unchanged whether or not in active use. For 7-person lab (1 PI + 6 FTE) the average lab size was 111.5 m2 (1,200 Net Assignable Square Feet). Using the surface and the average reported energy consumption per m^2^ we could estimate the footprint per study^20^. We estimate that 73% of the footprint is assignable to electricity demand and 28% to heating/cooling of the building (Figure 1C). Of note, this estimation accounts for the demand from equipment in use within the lab (15% of the total energy demand), the ventilation (35% of the total) and heating of the lab (28% of the total energy demand). For all studies, the energy demand amounted to 96.9 GWh (assuming 1 year of study completion time). This is close to twice as high as the reported consumption of the European Molecular Biology Laboratory (EMBL; 1,800 FTE; 57.8GWh) for 2021 alone, which has less than 100% laboratory space^21^. During the period from 2010 to 2024, we estimated a total consumption of 63.8 GWh amounting to 21,590 tons of CO2e. Laboratory energy demand and its associated footprint was the largest of all sectors we used in our estimation (Figure 1B).

Taken together, we estimated that the studies published from 2010 to 2024 generated 30,068 tons of CO2e. This is equivalent to 90 million km driven with a medium-size car (for comparison, the moon is only 0.3844 million km away), or above the footprint of the entire University of Bristol in a year (24,925 tons of CO2e scopes 1 and 2 for 2022/23)^22^. Our estimates indicate a large CO2e footprint associated with the failure of this literature to self-correct from 2010.

## DISCUSSION

Our environmental impact assessment of the failure of the 5-HTTLPR field to self-correct highlights the importance of self-correction in scientific research. This should in principle be achieved by swiftly correcting erroneous findings when new well-supported evidence is produced. The aim of our work was not to blame individual researchers, who are merely working within existing structures that incentivise publication or may not be aware of contradictory findings, but rather to utilise known literature to provide a basis of estimating the environmental costs of potentially unnecessary research. We selected the 5-HTTLPR literature only because it presented a defined body of findings that were subsequently shown to be false, used common methodologies, and where evidence for the lack of a meaningful association was evident as early as 2005. Our message is not that the authors of those papers should be further scrutinised for resource consumption, but rather that, moving forward, science needs to recognise its impacts beyond publication and be better equipped to respond to results which challenge existing findings. Excluded from our assessment were estimated financial costs, which similarly to estimating emissions can be variable and challenging^23^, though we can assume that with the volume of publications assessed the sum would be significant.

One of the challenges faced by our study is the lack of data regarding the carbon footprint of various aspects of scientific operations, which would have enabled more accurate estimates of the environmental impacts of studies. We therefore relied on several assumptions. For example, we based the laboratory facility efficiency on energy figures from UK facilities, whereas some of the studies performed their laboratory work outside of the UK in facilities which may have different levels of environmental efficiency^10^. Wherever there was uncertainty, we avoided over-estimations. As a result, one could expect our overall estimates to be conservative. As such, our methodology for assessing retrospective costs and impacts is not the primary focus of this study, but rather highlights the high resource demand of research that fails to self-correct^24^. Other fields of research may require more resources than genotyping, which requires relatively small volumes of consumables, and would require new but likely comparable methods to assess. As more life cycle assessments (LCAs) of consumables^25^ and specialist research fields are conducted^26–28^, as well as more open-source emission calculators are developed^29,30^, the accuracy of future analyses in this field is poised to significantly improve^31^.

The output of scientific research increases every year at an annual rate of between 4% and 9%^32,33^. While this generates important advances in knowledge (and in turn health and technology), it ultimately translates into an increase in environmental footprint. Efforts to address the environmental sustainability of science have focused largely on the energy consumption of facilities and equipment, consumables and chemicals. What has been lacking from this discussion is whether much of the research conducted is necessary despite evidence from other sources highlighting the problems of research waste^24^. Our work highlights that the environmental impact of unnecessary research may be as important – if not more – than the operational efficiency of science facilities, and will require a different approach to address. Not every research avenue represents as much unnecessary research as the example we have selected, but it does underscore that sustainable science efforts must consider the necessity and quality of research^34^.

Solutions to the challenges we highlight will require a collaborative and coordinated effort across the research sector. Researchers must scrutinise literature prior to embarking on new studies to ensure that foundational work is robust. Citation biases can provide an unfounded sense of robustness, so that ideally this would be supported by a systematic review of new literature, particularly where there are findings which challenge existing dogma. Perhaps more importantly, funders, publishers and institutions must work together to incentivise publication quality over quantity, and ensure practices necessary for scientific self-correction are supported (e.g., publishing null results^31^, halting lines of research that are not justified, improving open access of research which may show conflicting results, validation studies etc.). Much commendable work has already commenced in this area, as the FAIR Guiding Principles outline^35^, as well as a number of research papers and funder and publication guidelines^36–39^.

In the face of the growing climate crisis and global health challenges, we increasingly rely on scientific research to show a path to a more sustainable society. We accept that conducting research can be resource-intensive, and we can accept this due to the positive impacts we associate with successful research. However, with the continued growth of science, and its resource-intensive nature, there is ever increasing responsibility to ensure our findings are accurate. Science has an obligation to meet the standards it promotes, and to ensure that it is self-correcting. As such, the scientific community must continue to reflect inwardly to ensure that results are as impactful as today’s global challenges necessitate.

## Methods

### Determining the point in time, and total studies, researchers, and patients

PubMed abstract and article information for papers was searched on 10^th^ July 2024 for mentions of “5-HTT AND depression” or “5-HTT AND anxiety” in title or abstract. Review articles were excluded from the list, leaving a total of 1,183 articles. To assess papers published after a reasonable cut off of 2010 (1 year post publication of the second large study, and 5 years post publication of the first large study showing the lack of relation between anxiety traits and 5-HTTLPR), we selected PubMed records of studies published from 2010 (n= 779 out of 1,183).

The PubMed records were analysed using custom R scripts (https://github.com/BenNicolet/irreproducibility-footprint). We obtained the total number of authors in the records by searching all studies and summing up all authors listed (n= 8,805 in 1,183 studies (1996-2024), and n= 5,863 in 779 studies (2010-2024)). This is comparable to the 455 studies found to review depression in 5-HTTLPR polymorphisms between 1991-2016 by Border et al^8^, though differs due to differences in how studies were identified. The median number of authors per article was 7 for either 1996-2024 or 2010-2024 periods, and we assumed to be 1 principal investigator (PI) + 6 full-time equivalent researchers (1 PI + 6 FTE).

To estimate the total number of patients assessed in these studies, we used sample numbers of Karg and colleagues^40^ which reported 755 samples per study. As a result, we estimated a total of 893,165 patients studied. Note that we did not include estimates on discarded assays. It is likely that due to technical problem in genotyping, or failed assay, not all assays are reported in studies, and that the overall sample number is greater than our estimation.

### Assessing carbon emissions

While the publications in our PubMed records spanned over two decades, we decided to focus on a single year for our carbon emissions assessment. We reasoned that the average year should be used (2011; 1996 to 2024). Carbon factors for 2011 were used for our calculations, or to the year closest were not available.

### Genotyping

We screened the methods section of the research papers in our PubMed records to isolate details on the genotyping procedure. We identified a methods detailing a genotyping protocol from 2007^41^, the same year we used for carbon factors. We used this as a reference to list the common consumables necessary for each assay. Each consumable was assessed for weight, and then converted into CO_2_e by utilising Defra’s carbon factors for plastics (first available figures: 2012). Notably we omitted the travel associated with patients commuting to the point of sampling, or the sampling of foreign patient populations outside of the researcher’s country.

### Commuting of researchers

To assess the carbon impact of commuting, we used published data on the average commuting distance^42^ and the average CO2e per km of distance travelled^43^. We estimated that each researcher commuted 5 days per week, for 52 weeks per year excluding 5 weeks of annual leave per annum. We computed the impact of the study duration by varying the completion time of each study from 6 months to 3 years. For the total footprint estimation, we assumed a completion time of 1 year per study.

### Travel to conference

To estimate the CO2e of traveling to conferences, we used published conferencing data from Ciers and colleagues^17^, which provided the estimated average carbon emissions for various scientific positions per annum, including PhD students, senior scientists, postdocs, and professors. We applied their distribution of posts to our author list, and thus generated total travel emissions per scientist.

### Energy consumption of laboratory spaces

To obtain an estimate of lab space used by labs, we used the median number of authors per study (7; 1PI + 6 FTE) and the average laboratory space occupied by 1 PI + 7 FTE (111.5m^2^)^44^. We used energy consumption data from the UK’s S-Lab laboratory energy audits which provided an average energy consumption per square meter^20^, and obtained the CO2 equivalent footprint using Defra’s 2011 emissions data.

Data and scripts are available on github: https://github.com/BenNicolet/irreproducibility-footprint

## ACKNOWLEDGEMENTS

We would like to thank Professor Kate Tilling and Professor Ton Schumacher for their critical comments on this manuscript.

## Author contribution

MF, BPN and MM conceived the study. PubMed data analysis conducted by BPN. All carbon emission estimate was conducted by MF and BPN, with support from AL. All figures were created by BPN.

## Data availability

All scripts, computer codes and analysis outputs are available on GitHub (https://github.com/BenNicolet/irreproducibility-footprint).

---

**Figure S1.**
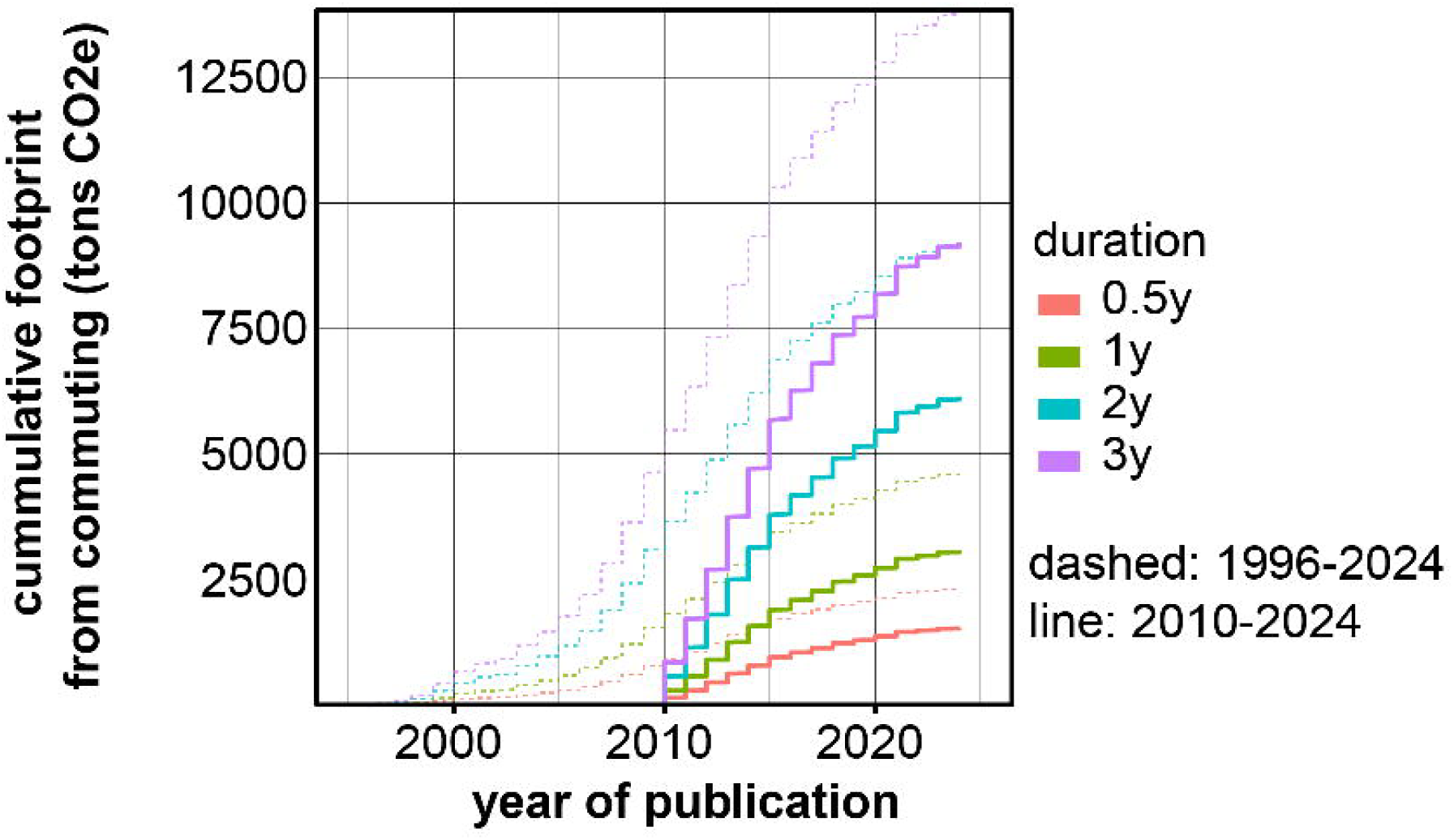

